# Antimicrobial Evaluation of *Salvadora persica* Methanolic Stem Extract and *In vivo* Modelling of its Identified Constituting Compounds Revealed Antibacterial Effect of 2-[(5-Iodosalicylidene)hydrazino]-4-morpholino-6-(1-pyrrolidinyl)-1,3,5-triazine in the Treatment of Dental Plague

**DOI:** 10.1101/2023.06.25.546444

**Authors:** Temidayo Olamide Adigun, Maikarfi Muhammed, Adisa Mofoluwaso Funmilayo, Abdulhameed Oluwatomi Alli, Fagboro Oluwayemisi Olufunke, Oyebamiji Samson Oluwatosin, Akinsehinwa Samuel Ayodeji, Aina Kehinde Oyebola

**Author notes:** Corresponding author (Adigun Temidayo Olamide).

## Abstract

Challenges of resistance to synthetic antimicrobials have opened new vistas in the search for natural products. This research rigorously reviews plant used in oral health. The antimicrobial activities of *Salvadora persica* commonly used as chewing stick was investigated against 3 clinical strains of *Escherichia coli, Staphylococcus aureus, Aspergillus nigar*. The antibacterial and antifungal activities of the extracts were determined using the agar well diffusion. *Salvadora persica*, was active against all the isolates especially on the bacteria with a MIC and MBC of 12.5mg/mL and 25mg/mL respectively. Phytochemical screening revealed the presence of alkaloids, tannins, flavonoids, saponins and traces of terpenoids. *In silico* modelling of the compounds identified in the chewing stick extract reveals that the aromatic nitrogen-rich compound with PubChem ID: 135580681 is likely responsible for the mechanistic inhibition associated with antibacterial effect of the plant material against dental plagues

## Introduction

Dental plaques (DPs) are complex biofilm found on salivary tooth surfaces. The pathogenesis of DP depends on factors such as adhesion of bacteria to salivary components, and adsorption to the tooth surface [1]. Gingivitis is a plaque-induced, and most common periodontal disease, and it results from activities of biofilm-forming bacteria located at the gingival margin. Gingivitis is the inflammation of gingival tissue, and its symptoms ranges from redness to swelling and bleeding of the gums. An untreated or poorly treated gingivitis may lead to loss of bone and tissue decay. This occurs via the combined activities of microorganisms within the subgingival biofilms and the host responses to them, lead to the progression of the disease and tissue damage [1]. By causing decalcification and eventual decay, Periodontopathogens such as *Porphyromonas gingivalis, Prevotella intermedia, Tannerella forsythia, Aggregatibacter actinomycetemcomitans, Fusobacterium nucleatum*, and *Capnocytophaga* sp., *Streptococcus faecalis* and *Streptococcus mutans*, have all been implicated in the pathogenesis of dental caries [1]. Plants are used as cosmetics, food, flavors, ornamental, and medicine. Due to their potential health benefits, medicinal plants have become crucial part of alternative medicine across the globe [2]. *Salvadora* (toothbrush tree/ *miswak*), belonging to the Salvadoraceae family, is one of the most important ones among 182 species of plants being used as chewing sticks. The roots, twigs, and stems of this plant have been used for oral hygiene in many Asian, Middle Eastern and African countries [3]. It has been reported that the aqueous and methanol extract of *miswak* has biochemical activity against organisms involved in the pathogenesis of DP [4]. Palombo [5] and Omer *et al*. [6] have also reported that methanolic extract of *S. persica* reduces adhesion of microbial pathogens to the tooth surface, which is a main event in the formation of DP and the progression to tooth decay and periodontal diseases.

Therefore, the aim of this research was to examine the potential of methanolic extract of *Salvadora persica* to inhibit bacterial and fungal growth, and thus, develop a strategy against formation of dental biofilms which may proffer viable and cost-effective alternative (s) to current remedy for dental plagues.

## 2.0 Material and Methods

### 2.1 Sample Collection and Preparation

Dried stems of *Salvadora persica* (*aswaki*) were bought from local Yoruba and Hausa herb sellers at the Sheik Gumi Market (Central Market) Kaduna State, Nigeria. The samples were identified and authenticated by Prof. Mahmoud Bala of Department of Plant Biology (Botany), Ahmedu Bello University (ABU) Kaduna State, Nigeria. The chewing sticks samples were washed under running tap water to remove dirt. The samples were air-dried for 2 days to curb distortion in the composition of the active principle in the chewing sticks. The dried samples were well pulverized into a fine powder with a mixer grinder. The powder was stored in air tight aseptic containers (28ºC±2) for subsequent use.

### 2.2 Preparation of Extracts

The Methanolic extracts were prepared using the maceration method of Adekunle and Odukoya [7] with slight modifications. The dried stem was crushed into fine powder. About 60g of the powder were separately soaked in 180ml of 95% methanol in a 250ml reagent bottle and stoppered. This was allowed to stand for 7 days in a dark well tighten bottle with frequent shaking to permit full extraction of the active principles in the powdered chewing sticks. The fluids were then filtered using Whatman No1 filter paper. The plant macerate was evaporated to dryness under reduced pressure at 40C. It was then kept in the fridge (4ºC) prior.

### 2.3 Phytochemical Analysis of Chewing Stick Extracts

Qualitative screening of the phytochemical components of the chewing sticks was carried out using the method outlined by Harborne [8] to detect the presence of glycosides, alkaloids, saponin, tannins, flavonoids anthocyanin, anthraquinone and phlobatannin.

### 2.4 Preparation of Media and Test Organisms

The method of Habamu *et al*. [9] was used for media preparation, and the clinical isolates of *staphylococcus aureus, Aspergillus niger*, and *Escherichia coli, were* obtained from the microbiology laboratory of Kaduna polytechnic, applied biology department

#### 2.4.1 Authentication of Test Organisms and Preparation of Nutrient Broth

Isolates of *staphylococcus aureus, Escherichia coli*, and *Aspergillus niger* were used for the study, and they were characterized by observing their cultural characteristics when sub-cultured on nutrient agar and the gram staining reaction for bacteria. Biochemical tests for motility, citrate, and methyl red as positive confirmatory tests were carried out for the test organisms. The nutrient broth was prepared using the method described by Habamu *et al*. [9].

### 2.5 Preparation of Different Concentration of Extracts

The method described by Perez *et al*. [10] was used for preparation of different extract concentration. Briefly, different concentration of extract was prepared (i.e 1000 mg/ml, 50 mg/ml, 25 mg/ml, and 12 mg/ml) since 1g equals 1000 mg, to concentration of 100 mg/ml, 1 g of each extract was dissolved in 10ml sterile distilled water. This was also used as stock solution from which the remaining concentration were prepared and made up to 10ml and each concentration was labelled.

#### 2.5.1 Preparation and Standardization of Inoculum

The overnight culture of nutrient broth of each bacteria isolate which was previously prepared was used to prepare inocula by diluting with sterile saline. Using the method described, Habamu *et al*. [9], 10ml of the solution was transferred into three (3) clean separate test tube sand were autoclaved at 121°C for 15min. the sterile saline was allowed to cool and 0.1ml of the overnight broth culture of *Escherichia coli, Staphylococcus aureus* and *Aspergillus niger*. Were dispensed into separate test tube containing the sterile normal saline. These served as standard inoculum which was used for the antibacterial and antifungal testing.

### 2.6 Determination of Antibacterial and Antifungal Activity Using Agar Well Diffusion Method

The agar well diffusion method of Habamu *et al*. [9] as described, was adopted to test for the antimicrobial activity of the extracts on the test organisms. 15cm sterile disposable petri dishes of nutrient agar were prepared and allowed to solidify. These plates were then separately flooded with diluted standardized overnight cultures by a loopful of bacteria/fungi inocula taken and streaked over the entire surface of the dried agar. Wells of 6mm diameter were made in triplicates in each plate with 0.1ml of diluted concentrations (100mg/ml, 50mg/ml, and 25mg/ml) of extracts with the aid of sterile pipettes per well. While 1mg/ml of the standard antibiotics’ ciprofloxacin were used as positive control. Sterile distilled water was used as negative control on a separate plant. Diameters of zones of inhibition were measured using millimeter rule after incubating the plates at 37°c for 24hrs. The plates were replicated in triplicates and the means of the zones of inhibitions for each organism at each concentration of extract were calculated and recorded (Perez *et al*. [10]).

### 2.7 Determination of Minimum Inhibitory Concentration (MIC) and Minimum Bactericidal-fungicidal Concentration

The minimum inhibitory concentration was carried out for each organism. Double strength of nutrient broth was prepared. Then 1ml of double strength was added into four test tubes for each organism and extract labelled 100mg/ml, 50mg/ml, 25mg/ml, and 12.5mg/ml. equal volume of the above concentration were incorporated in nutrient broth in 1:1 ratio and 0.1ml of standard suspension of test tube organisms and was added to each of the test tube were then incubated aerobically at 37°C for 24hours and anaerobically at 25-27°C (ambient temperature) for fungi. Tubes containing broth and extract without inoculum were included to serve as positive control while a tube containing broth and inoculum serves as negative control for comparison. The presence of growth (turbidity solution) or visible growth was recorded as the minimal inhibitory concentration. Minimum Bactericidal/fungicidal Concentration was determined using the method of Vinothilkumar *et al*. [11].

### 2.8 *In-silico* Modelling of *Salvadora persica* Compounds

The crystal structures of DNA Gyrase subunit A and B (PDB ID: 3NUH) were retrieved from RCSB Protein Databank (http://www.rcsb.org). The various co-crystal ligands and water molecules were deleted from the original protein crystal. The 2D structures of the identified compounds, through gas chromatography-mass spectrometry, from *Salvadora persica* extract, as well as levofloxacin (standard inhibitor) were downloaded from the PubChem database and prepared in a pdbqt file with appropriate torsion numbers. Molecular docking of the compounds was carried out using Autodock vina [17]. The Root Mean Square Deviation (RMSD) and Affinity Energy [17] were used in selecting the best interaction poses. The protein-ligand interactions were examined and represented using Discovery Studio Visualizer version 21.1.0.20298 and Schrodinger PyMol® (TM) version 2.5.1 (version 16).

## 3.0 Results

### 3.1 Extraction Yield, Qualitative and Quantitative Phytochemical Screening

The extraction of *Salvadora persica* using methanol as solvent yielded 20 %. Table 3.1 revealed the presence of phytochemicals such as anthraquinones, flavonoids, terpenoids, saponins, alkaloids, tannins, phenols, and cardiac glycosides.

**Table 3.1.** :Phytochemical Composition of Methanolic Stem Extract of *Salvadora persica*.

| Constituents | Results |
| --- | --- |
| Flavonoids | +ve |
| Saponins | +ve |
| Cardiac glycosides | +ve |
| Alkaloids | +ve |
| Anthraquinone | +ve |
| Tannins | +ve |
| Terpenoids | +ve |
| Phenols | +ve |

### 3.2 Antimicrobial Activity of Salvadora Extract Against the Test Organisms

The extract shows high activity on *staphylococcus aureus* having zones of inhibition 22.1 at 100mg/ml, *Escherichia coli* 15.4 at 100mg/ml, and *Aspergillus niger* 18.6 at 100mg/ml concentration. At least concentration *staphylococcus aureus* shows 11.0 at 12.5mg/ml, *Escherichia coli* 8.4 at 12.5mg/ml, and *Aspergillus niger* 6.2 were inhibited (Table 3.2).

**Table 3.2.**
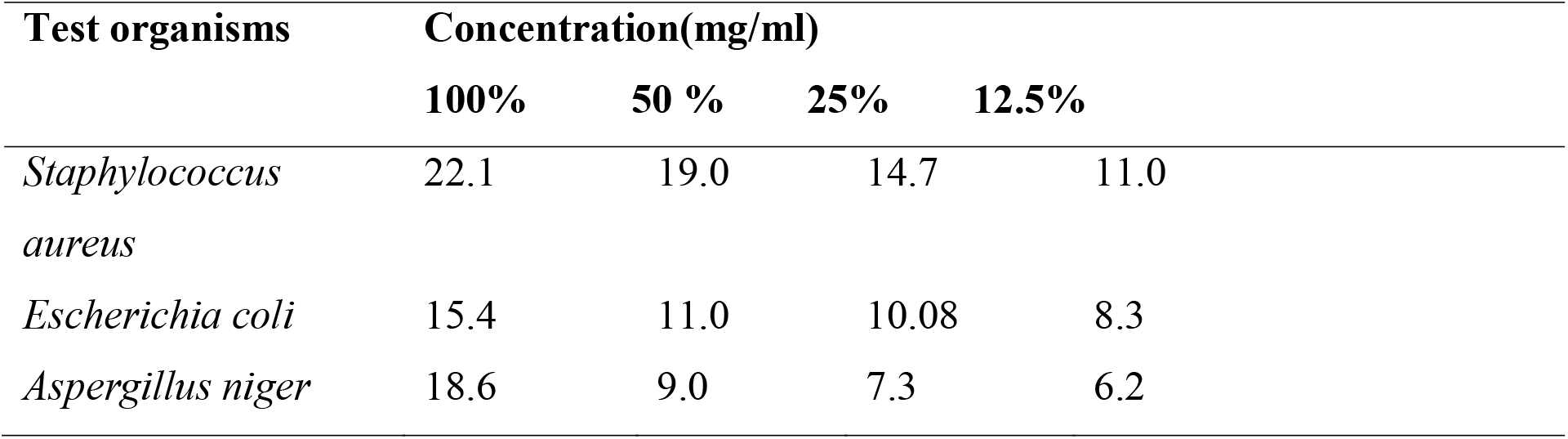
:Antimicrobial Activity of Salvadora Extract Against the Test Organisms.

### 3.4 The Minimum Fungicidal Concentration (MFC) of *Salvadora persica* Methanolic Stem Extract Against the Fungi Isolate

Methanolic extract of *Salvadora persica* showed a high antifungal activity as shown in Table 3.4. A significantly higher antibacterial activity was exhibited by methanol extract and *Salvadora persica*. The control set up did not show any antimicrobial activity.

**Table 3.3.** :The Minimum bactericidal concentration (MBC) of *Salvadora persica* Methanolic Stem Extract Against the Bacteria Isolate. All concentrations except 100 showed turbidity (table 3.3)

| Bacteria isolates | Concentration (m/ml) |  |  |  |
| --- | --- | --- | --- | --- |
|  | 100 | 50 | 25 | 12.5 |
| *Staphylococcus aureus* | - | + | + | + |
|  | - |  |  |  |
| *Escherichia coli* | - | + | + | + |

**Table 3.4.** :The Minimum Fungicidal Concentration (MFC) of *Salvadora persica* Methanolic Stem Extract Against the Fungi Isolate.

| Fungi isolates | Concentration (mg/ml) |  |  |  |
| --- | --- | --- | --- | --- |
|  | 100 | 50 | 25 | 12.5 |
| *Aspergillus niger* | - | - | + | + |

### 3.5 The Minimum Inhibitory Concentration (MIC) of *Salvadora persica* Methanolic Stem Extract Against the Bacteria Isolate

At 50 and 100 mg/ml, the extract showed high minimum inhibitory concentration (Table 3.5).

**Table 3.5.** :The Minimum Inhibitory Concentration (MIC) of *Salvadora persica* Methanolic Stem Extract Against the Bacteria Isolate.

| Bacteria isolates | Concentration (mg/ml) |  |  |  |
| --- | --- | --- | --- | --- |
|  | 100 | 50 | 25 | 12.5 |
| *Staphylococcus aureus* | - | - | + | + |
| *Escherichia coli* | - | - | + | + |

### 3.6 The Minimum Inhibitory Concentration (MIC) of *Salvadora persica* Methanolic Stem Extract Against the Fungi Isolate

**Table 3.6.** :The Minimum Inhibitory Concentration (MIC) of *Salvadora persica* Methanolic Stem Extract Against the Fungi Isolate. All concentrations except 12.5 mg/ml (table 3.6).

| Fungi isolates | Concentration (mg/ml) |  |  |  |
| --- | --- | --- | --- | --- |
|  | 100 | 50 | 25 | 12.5 |
| *Aspergillus niger* | - | - | - | + |
| Key: + = Growth observed |  | - = No Growth observed |  |  |

### 3.7 Binding Affinity Results of Modelled Compounds of *Salvadora persica* Methanolic Stem Extract Against DNA Gyrase Subunit A, as well as their Interactions

#### 3.7.1 Binding Affinity of Modelled Compounds of *Salvadora persica* Methanolic Stem Extract Against DNA Gyrase Subunit A

The compound with PubChem ID: 135580681, humulene, caryophyllene, beta-Selinene, as well as 899374-61-9 exhibited comparatively higher binding affinity than levofloxacin (standard inhibitor) (−6.1 Kcal/mol) on DNA gyrase subunit A, with compound with PubChem ID: 135580681 (−8.0 Kcal/mol) displaying the highest binding affinity value (Figure 1).

**Figure 1:**
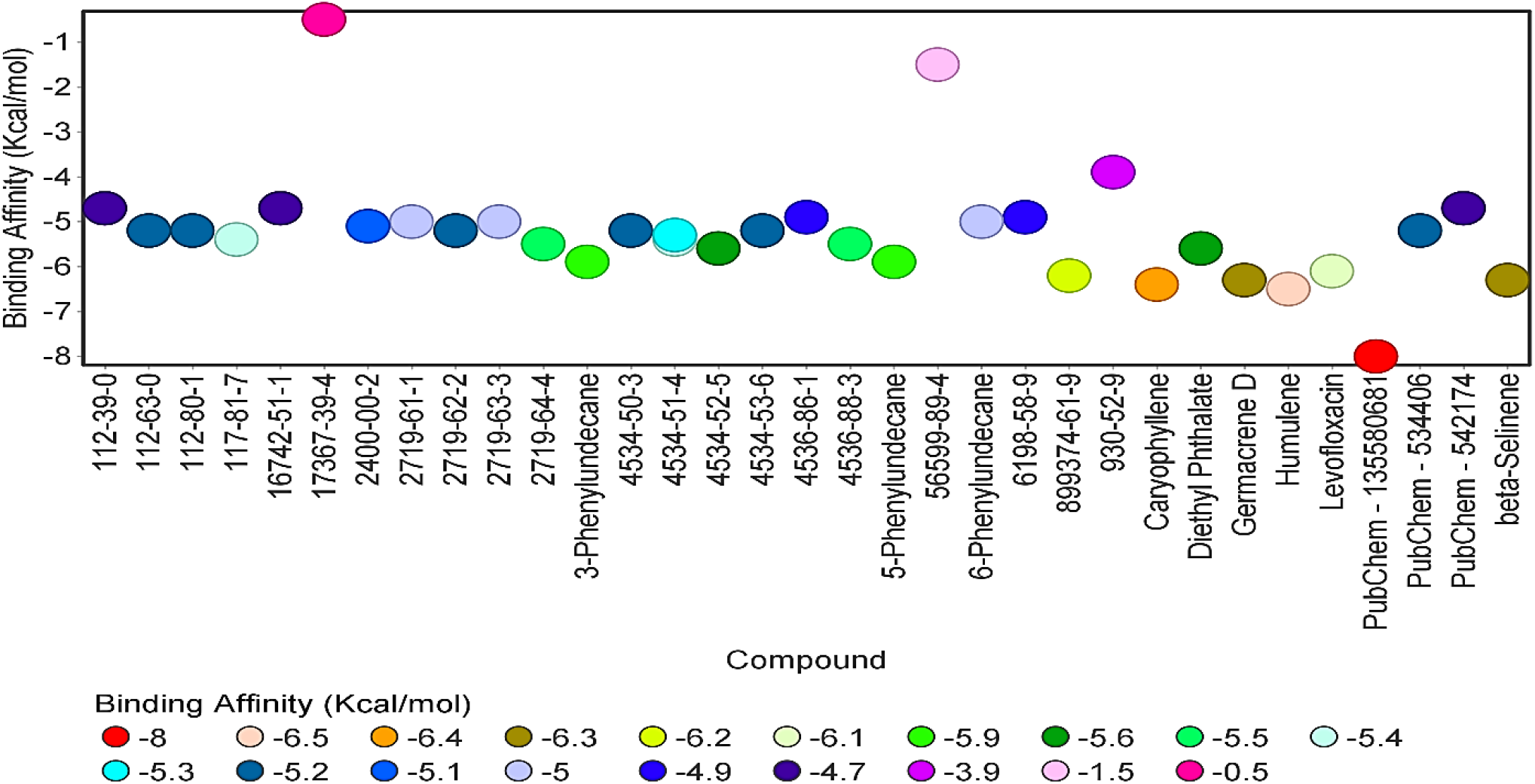
Binding Affinity Results of Identified Compounds of *Salvadora persica* Methanolic Stem Extract Against DNA Gyrase Subunit A.

#### 3.7.2 Comparative Interactions of PubChem – 135580681 (Hit Compound) and Levofloxacin (Standard Inhibitor) on DNA Gyrase Subunit A

Levofloxacin binds with tryptophan 59 (at a distance of 2.41 Å) and aspartic acid 137 (at a distance of 2.37 Å) of DNA gyrase subunit A pocket through the conventional hydrogen bonding; alanine 136 through carbon-hydrogen bonding (at a distance of 3.45 Å); glutamic acid 139 (at a distance of 3.43 Å) through the ionic halogen bonding, as well as the active hydrophobic histidine 132 through the hydrophobic Pi-cation bond (at a distance of 4.78 Å) (Figure 2).

**Figure 2:**
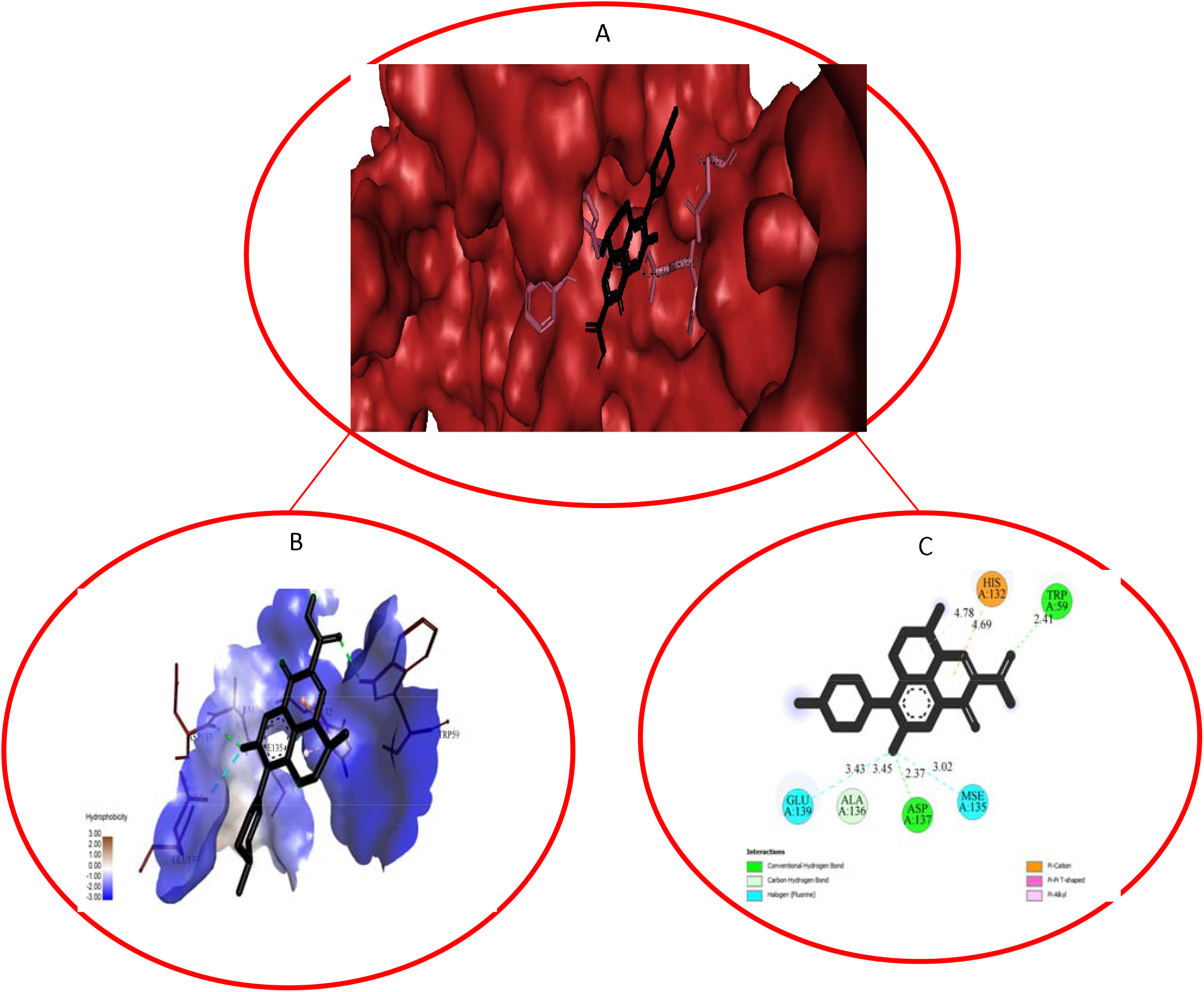
Pocket View (A), Pocket Hydrophobicity View (B), and Two-Dimensional (2D) View (C) of Interaction Between Levofloxacin and DNA Gyrase Subunit A.

The compound with PubChem ID: 135580681 binds with histidine 132 through carbon-hydrogen bond (at a distance of 3.52 Å); active asparagine 53 (at a distance of 2.54 Å); alanine 136 through the hydrophobic Pi-alkyl bond (at an average distance of 7.11 Å) which are in common with the binding pattern of levofloxacin, in addition to binding to side residues leucine 138 (at a distance of 3.17 Å), as well as tyrosine 50 through the hydrophobic Pi-Pi T-shaped interactions (at a distance of 5.37 Å) (Figure 3).

**Figure 3:**
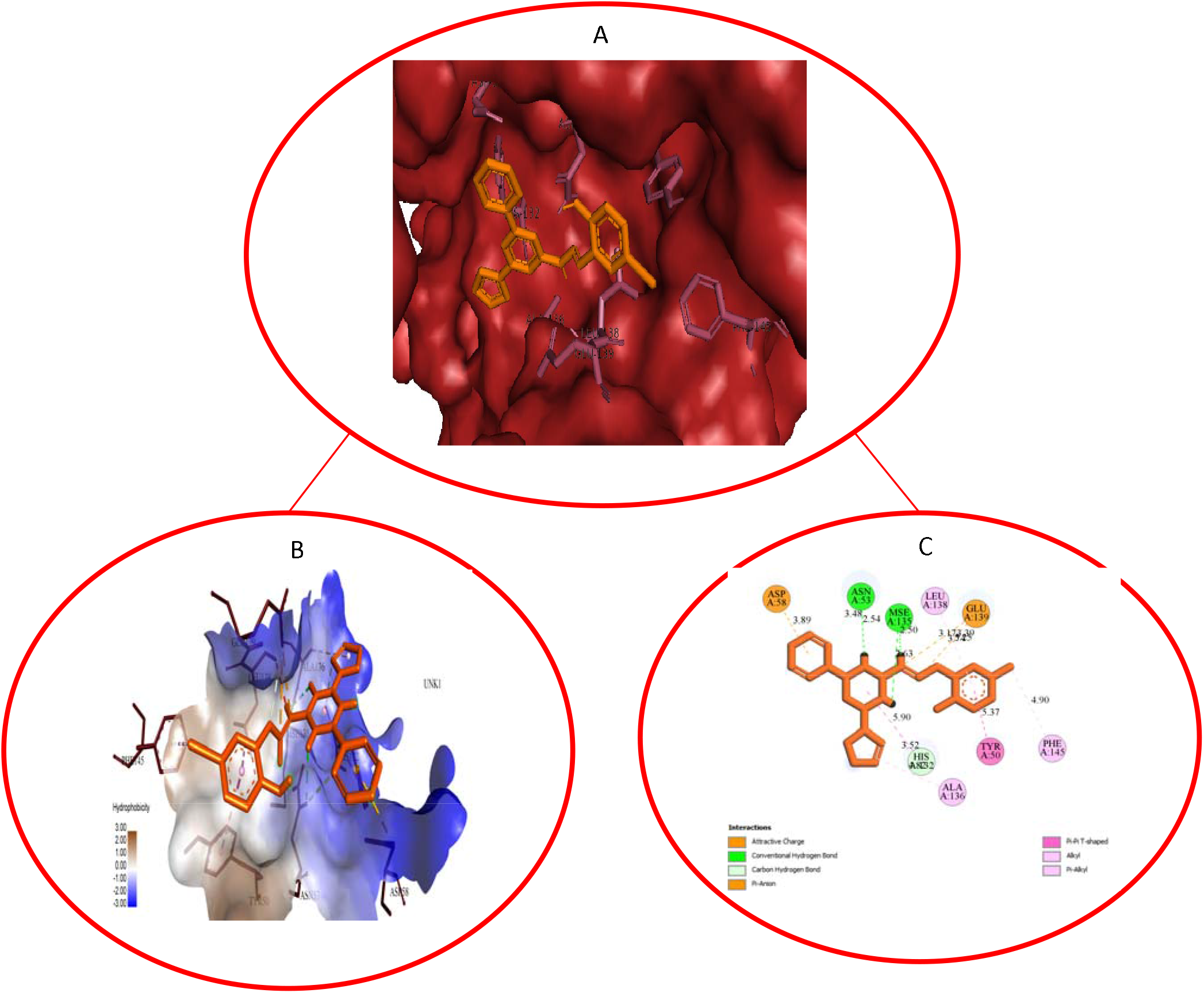
Pocket View (A), Pocket Hydrophobicity View (B), and Two-Dimensional (2D) View (C) of Interaction Between PubChem - 135580681 and DNA Gyrase Subunit A.

**Figure 4:**
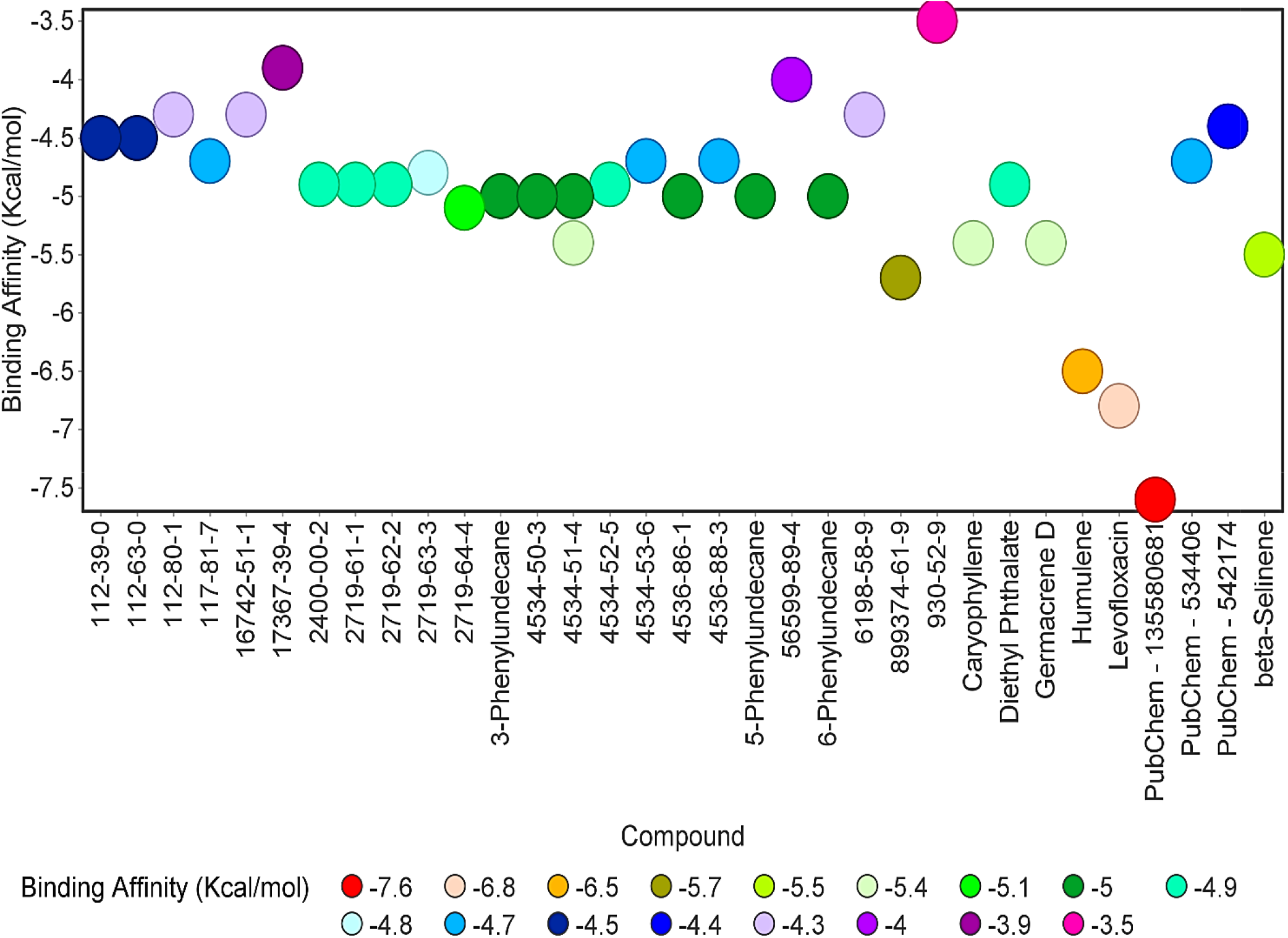
Binding Affinity Results of Identified Compounds of *Salvadora persica* Methanolic Stem Extract Against DNA Gyrase Subunit B.

3.8 Binding Affinity Results of Modelled Compounds of *Salvadora persica* Methanolic Stem Extract Against *Escherichia coli* K-12 DNA Gyrase Subunit B, as well as their Interactions

3.8.1 Binding Affinities of Modelled Compounds of *Salvadora persica* Methanolic Stem Extract Against DNA Gyrase Subunit B

All the compounds exhibited comparatively higher binding affinity than levofloxacin (standard inhibitor) (−6.8 Kcal/mol) on DNA gyrase subunit A, except the compound with PubChem ID: 135580681 (−7.6 Kcal/mol) which displayed comparatively higher binding affinity value than the standard.

#### 3.8.2 Comparative Interactions of PubChem – 135580681 (Hit Compound) and Levofloxacin (Standard Inhibitor) on DNA Gyrase Subunit B

Levofloxacin binds with leucine 534 and alanine 499 through the alky and Pi-sigma bond at the distance of 3.50 Å and 3.74 Å respectively; binds to proline 532 and lysine 740 through carbon-hydrogen bond at the distance of 4.02 Å and 4.56 Å, as well as the hydrophobic arginine 738 through Pi-cation interaction at the distance of 4.16 Å (Figure 5).

**Figure 5:**
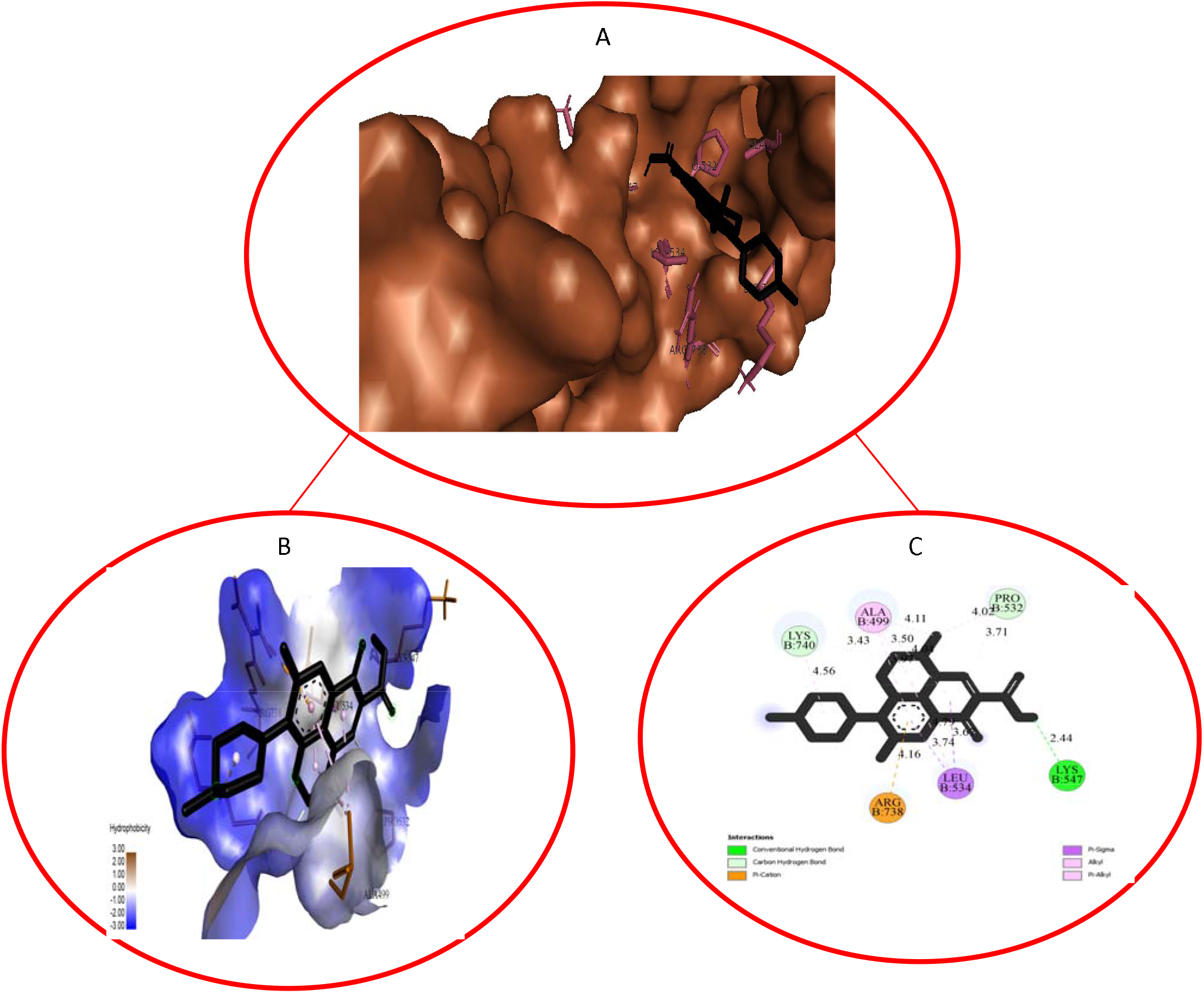
Pocket View (A), Pocket Hydrophobicity View (B), and Two-Dimensional (2D) View (C) of Interaction Between Levofloxacin and DNA Gyrase Subunit B.

Compound with PubChem ID: 135580681 binds (in common with levofloxacin) to leucine 534 and alanine 499 through alkyl bond at the distance of 4.28 Å and 4.64 Å respectively; proline 532 through Pi-alkyl interaction (at a distance of 4.47 Å); active aspartic acid 548 through the ionic salt bridge (at an average distance of 2.83 Å), in addition to the interaction with the side chain residue glutamine 531 through carbon-hydrogen bond (at a distance of 2.66 Å). (Figure 6).

**Figure 6:**
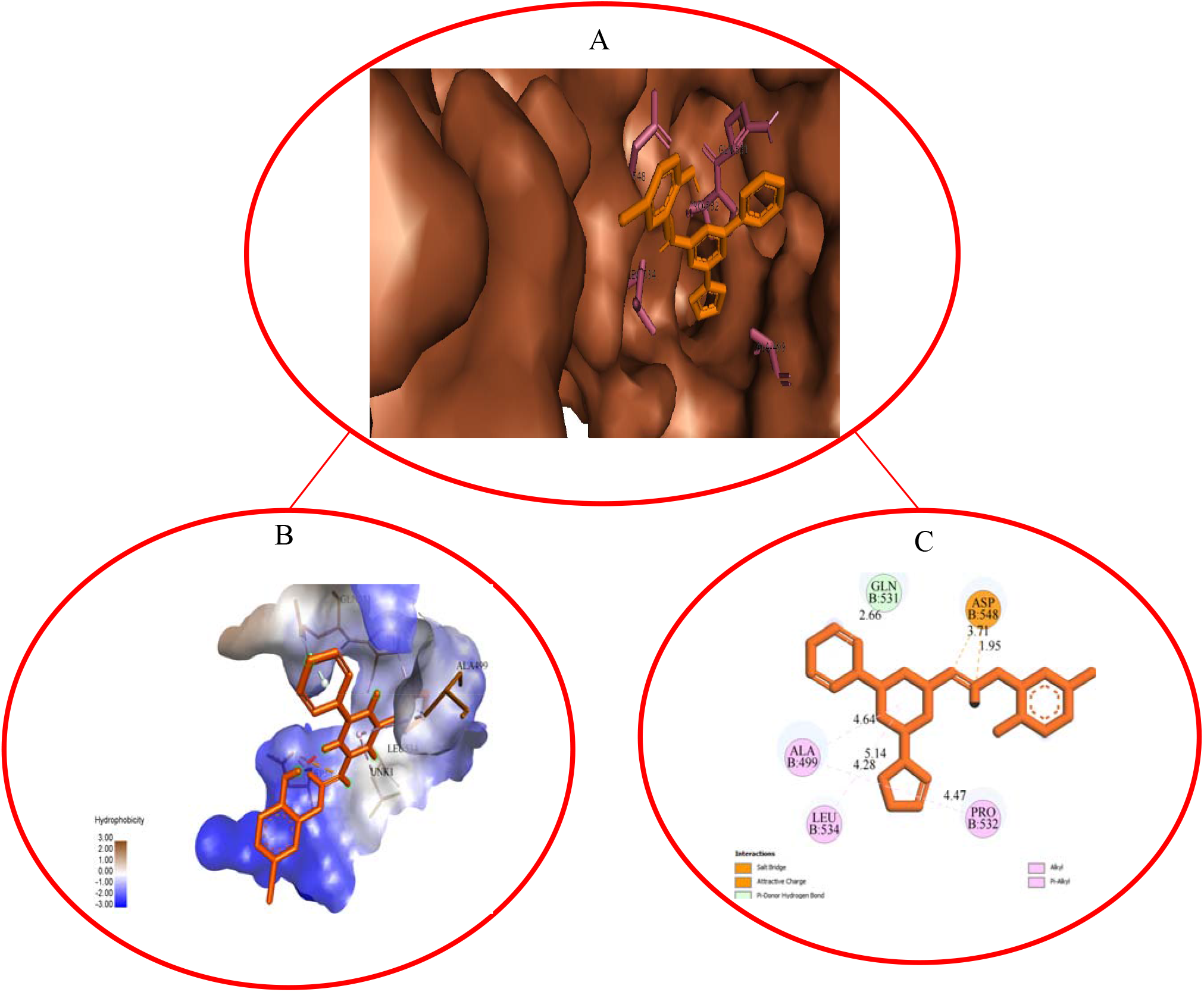
Pocket View (A), Pocket Hydrophobicity View (B), and Two-Dimensional (2D) View (C) of Interaction Between PubChem – 135580681 and DNA Gyrase Subunit B.

## 4.0 Discussion

*Salvadora persica* chewing stick has been reported to possess a greater pharmaceutical effect which might be due to its bioactive components [7]. The present study showed that the extract of tropical Nigerian Chewing stick; *Salvadora persica* exhibited a great antifungal and antibacterial effects against the test organisms. *Salvadora persica* showed a high inhibitory activity against the bacteria species tested, since it exhibited the highest antibacterial inhibition. Results obtained showed that methanol was an effective and powerful solvent. This is consistent with the report of Kareem *et al*. [12] which stated that active components of plants are more soluble in organic solvent. The high potency of the methanol extract may be attributed to the dissolving power of alcohol [13]. The antifungal and antibacterial inhibitory effects showed by the chewing stick extract may be attributed to the presence of phytochemical compounds (glycosides, alkaloids, saponin, tannins, flavonoids anthocyanin and anthraquinone) present in them. This is also in agreement with the work reported by Cowan [14]. Antimicrobial potency of *Salvadora persica* has been the major reason why they are being employed in tropical Nigeria for dental cleaning to guide against dental caries, gingivitis, and dental plaque [15]. From the result obtained, it was observed that Minimum Inhibitory effect was vividly shown at varying concentrations (MIC) of extracts of the two chewing sticks. The extract shows high activity on *staphylococcus aureus* having zones of inhibition 22.1 at 100mg/ml, *Escherichia coli* 15.4 at 100mg/ml, and *Aspergillus niger* 18.6 at 100mg/ml concentration. At least concentration *staphylococcus aureus* shows 11.0 at 12.5mg/ml, *Escherichia coli* 8.4 at 12.5mg/ml, and *Aspergillus niger* 6.2 were inhibited.

The great antibacterial effect observed in this study showed by the chewing sticks was because of tannin present in them [15]. Hagerman and Butler [16] have reported that tannins have been shown to form irreversible complexes with proline-rich proteins which would lead to inhibition of cell wall-protein synthesis, a property that may explain the mode of action of these chewing stick extracts. *In silico* modelling of the compounds identified in the chewing stick extract reveals that the aromatic nitrogen-rich compound with PubChem ID: 135580681 is likely responsible for the mechanistic inhibition associated with antibacterial effect of the plant material against dental plagues observed in this study, with the mechanism of inhibition involving a mode similar to that of pan inhibitory activities of quinolone-derived levofloxacin against both the subunit A and B of the bacterial DNA gyrase. The structure name of the compound with PubChem ID: 135580681 is 2-[(5-Iodosalicylidene)hydrazino]-4-morpholino-6- (1-pyrrolidinyl)-1,3,5-triazine.

## Conclusion

This research, therefore, has revealed the antifungal and antibacterial effect of *Salvadora persica* methanolic stem extract in *in vivo* model of orodental hygiene, as well as the mechanistic inhibitory activity of its constituting 2-[(5-Iodosalicylidene)hydrazino]-4-morpholino-6-(1-pyrrolidinyl)-1,3,5-triazine against both the subunit A and B of *Escherichia coli* K-12 DNA gyrase (a major component microbial composition of dental plague) through *in silico* simulations. These findings also evidently support the view that chewing sticks may serve as an effective natural alternative source of antibacterial and antifungal agents against different oral diseases, along with additional interproximal cleaning aides. Toxicological studies are recommended to ascertain the toxicity status of the medicinal plant, and there is a need for further evaluation of 2-[(5-Iodosalicylidene)hydrazino]-4-morpholino-6-(1-pyrrolidinyl)-1,3,5-triazine in both pre-clinical and clinical studies for the development of natural, cost-effective, clinically safe, and highly potent anti-dental plague agent.

